# Antioxidant, anti-enzymatic, antimicrobial and cytotoxic properties of *Euphorbia tirucalli* L

**DOI:** 10.1101/2019.12.17.879692

**Authors:** Mohammad Altamimi, Nidal Jaradat, Saad Alham, Motasem Al-Masri, Ala Bsharat, Rania Alsaleh, Rozan Sabobeh

## Abstract

Medicinal properties of *Euphorbia tirucalli* L. have been investigated in vitro. Water extract from the plant latex was freeze dried and tested against 3 digestive enzymes and showed IC50 as 39.8±0.22, 79.43±0.38 and 316.22±0.3 for lipase, alpha- glucosidase and alpha- amylase, respectively. These results were incomparable with drugs such as acarbose and orlistat. Antioxidant property showed that the IC50 values of *E. tirucalli* and Trolox were 79.43±0.69 and 3.1 ±0.92 respectively. Also, the plant extract exhibited a range of antimicrobial activity against gram-positive and gram-negative bacteria and pathogenic yeast and fungi. Cytotoxicity of the plant extract tested on Caco-2 cells was determined using MTT method. The effect was linear and almost 0.5 mg/ml extract has inhibited 50% of cells relative to the control. *Euphorbia tirucalli* showed promising activities and is potential source of active ingredient with functional properties.

## Introduction

Phytogenic chemicals play an essential value in drugs investigations. Many of drug candidates and clinically approved therapeutic agents have been isolated from plant products [1]. In fact, several isolated molecules from herbal sources utilized in medicine, have the most potent therapeutic activity among chemically produced medicines such as vincristine, taxol, atropine, morphine, pilocarpine and many others [2]. The potent physiological and therapeutic effects of plant materials usually resulted from the effects of plant secondary metabolic compounds.

Diabetes mellitus, obesity, and overweight are dangerous metabolic disorders and considered the major reasons for mortality and morbidity for thousands of people in many of the developing countries. However, different investigations demonstrated that obesity and overweight are associated with an increased risk of diabetes. In fact, both of the obesity and overweight elevate the costs of immense health care because they are significantly associated with diabetes, arthritis, asthma, high level of blood cholesterol, high blood pressure and poor quality of life [3].

Malignant tumor is commonly known as cancer which is a large group of diseases that may affect every part of the human body with a rapid formation of abnormal cells and usually distributed to other organs. However, the death from cancer usually occurred from the metastases process and considered a second leading cause of death world widely. In 2018 cancer was responsible for about 9.6 million deaths with 70% of deaths were from middle- and low -income regions [4–6].

Evidently, the anti-microbial resistance is a global medical and public health concern as it is not limited to bacterial pathogens but also distributed to the parasites, fungi and viruses. Moreover, the most therapeutically effective and safe drugs have already been developed, and newer effective drugs often have more toxicity and other drawbacks, including higher cost [7].

The imbalance between antioxidants and free radicals in human body called oxidative stress. While oxidation is a normal internal biochemical process in living cells which results in production of free radicals causing long chains of chemical reactions with other compounds in the body, other external such as alcohol, high sugar, fat diet, pollution, radiation, smoking and pesticides are some of the risk factors of oxidative stress [8]. However, antioxidants compounds can donate an electron to free radicals sparing the reactions with vital molecules in the cell such as DNA or lipids of cell membrane. This can cause these free radicals to be more stable. In fact, oxidative stress can contribute to the development of many diseases including heart disease, inflammation, glaucoma, diabetes, chronic obstructive pulmonary disease, atherosclerosis and cancer [9].

*Euphorbia tirucalli* L. is a perennial laticiferous, evergreen shrubby plant belongs to Euphorbiaceae family. It grows wildly in tropical and subtropical regions or cultivated for decorative purposes as ornamental plant. The *E. tirucalli* branches have a pencil shape from which obtained its vernacular name, the pencil-tree plant. However, in all its parts, *E. tirucalli* contains white milky latex that contains triterpenoids as cyclotirucanenol [10, 11], diterpene tirucalicine [12], euphol, amyrin, cycloeuphordenol, tirucallol, lanosterol, glut-5-en-3-b-ol, cycloartenol euphorginol [13] and steroids [14].

In addition, *E. tirucalli* aqueous extract contains various types of phenolic acids and flavonoids including gallic acid, chlorogenic acid, caffeic acid, kaempferol, quercetin and rutin [15].

In various folk medicines, *E. tirucalli* white latex used for the treatment of warts, epilepsy, sexual impotence, snake bites, hemorrhoids, toothache, corn, rheumatism, acne [16], cancer [17] and antimicrobial agent [18, 19].

Some in vitro studies were found in the literature regarding *E. tirucalli* biological and pharmacological properties which revealed that the plant has antiviral, larvicidal, molluscicide and anti-arthritic activities [20–23]. Moreover, *E. tirucalli* showed cytotoxicity, genotoxicity and made some changes in antioxidant gene expression in human leukocytes [15].

On the other hand, African *E. tirucalli* latex revealed a co-carcinogenic property and specific cellular immunity reduction associated to the infection from Epstein-Barr virus [24].

The current investigation aims to evaluate the inhibitory effects of *E. tirucalli* freeze dried juice against α-amylase, α-glucosidase, lipase, microbial growth also aims to assess its antioxidant and cytotoxic properties.

## Material and methods

### Instrument for anticancer test

Microplate reader [Unilab, 6000, Mandaluyong, USA], CO_2_ incubator [Esco, 2012-74317, Changi, Singapore], inverted microscope [MRC, IX73, Hong Kong, China], UV-Visible Spectrophotometer [Jenway 7315, Staffordshire, UK], vortex [Heidolph Company, 090626691, Schwabach, Germany], ultrasonic cleaner [MRC Laboratory Equipment, 1108142200049, Essex, UK], autoclave [MRC Laboratory Equipment, A13182, Essex, UK], Water bath [Lab Tech,2011051806, S. Korea], Stir-Mixer [Tuttnaver, 300303159, USA], Cooled incubator [Gallenkamp, SG92/01/244, Loc, United Kingdom], Micropipette [MRC Laboratory Equipment, MPC-1000, Essex, UK], Multichannel Micropipette [MRC Laboratory Equipment, MPC-8-50, Essex, UK].

### Instruments for antioxidant, lipase, amylase and glycosidase inhibition tests

UV-Visible Spectrophotometer [Jenway 7135, England], filter papers [Whitman no.1, USA], shaker device [Memmert shaking incubator, Germany], rotatory evaporator [Heidolph vv2000 Heidolph OB2000, Germany], grinder [Moulinex model, Uno, China], balance [Red wag, AS 220/c/2, Poland] and freeze dryer [Mill rock technology BT85, China].

### Instruments for antimicrobial test

Sonicator [ultrasonic cleaner, MRC laboratory equipment, 1108142200049, Essex, UK], Autoclave [MRC laboratory equipment, A13182, Essex, UK], Water bath [Iso 9001 certified, Lab tech, 2011051806, Korea], Incubator [EN500, Nuve, A08 No.789244, Turkey], Stir-Mixer [vortex] [Tuttnaver co., 300303159, USA], Cooled incubator [Gallenkamp, SG92/01/244, Loc, UK], Micropipette [MRC, MPC-1000, UK], Multichannel Micropipette [MRC, MPC-8-50, UK].

### Chemicals and reagents

BBL Mueller Hinton II broth, cation adjusted [Becton Dickinson, USA], Difco Sabouraud agar, modified [Becton Dickinson, USA], Acarbose, pNPG, α-glucosidase [Baker’s Yeast alpha glucosidase], α-amylase, DNSA and potassium phosphate from Sigma-Aldrich, USA. Methanol, NaOH, n-hexane, and acetone were purchased from Lobachemie [India], [DPPH] 2,2-Diphenyl-1-picrylhydrazyl was obtained from Sigma-Aldrich [Germany] and DMSO [Dimethyl sulfoxide] was obtained from Riedeldehan [Germany]. In addition, Trolox [(s)-(-)-6 hydroxy-2, 5, 7, 8-tetramethychroman-2-carboxylic acid] was obtained from Sigma-Aldrich [Denmark]. Alpha-amylase [Sigma, Mumbai, India], DNSA [3, 5-dinitrosalicylic acid], Acarbose [Sigma, St. Louis, USA], p-nitrophenyl butyrate, Orlistat, tris-HCl buffer and Porcine pancreatic lipase type II were purchased from Sigma [USA].

### Plant material

The pencil shape branches of *E. tirucalli* were collected from Tulkarm region of Palestine in 2018. The taxonomical characterization was established at An-Najah National University in Natural Products laboratory, Department of Pharmacy and the herbarium was stored under the voucher specimen code Pharm-PCT-1002.

The collected branches were washed with sterile water and the cleaned branches were grated and pressed using mechanical Juicer Extractor Machine [Aicok Juicer, China]. Plant juice was sterilized utilizing Millipore Sigma membrane filtration device [Germany]. The produced liquid was dried using a freeze-drier apparatus and then kept in an air-tight brown jars at 4ºC for further use.

### Free radical scavenging assay

Methanolic stock solution [1mg/ml] was set for *E. tirucalli* dried juice and for vitamin E analogue [Trolox] which used as a control with a potent antioxidant activity. Then, various concentrations from the previous stock solution were prepared. Plant working solution [1 ml] was mixed with 1ml freshly prepared DPPH [0.002 g/ml] methanolic solution and 1ml methanol was then added to the previous mixture. The blank solution contained DPPH and methanol only in a ratio of 1:1. The solutions were incubated at room temperature [25ºC] in a dark place for 30 min. Then, their optical densities were measured by the UV/Vis spectrophotometer [Jenway 7135, England] at 517 nm.

Antioxidant activity was calculated per the following equation (1):

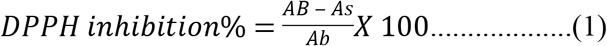

A_b_ is the recorded absorbance of the blank solution

A_S_ is the recorded absorbance of the sample solution or control.

### Porcine pancreatic lipase inhibitory assay

A porcine pancreatic lipase inhibition assay was carried out to assess the activity *E. tirucalli*. Orlistat, a commercially available anti-obesity, and an anti-lipase therapeutic agent was used as a reference control. The porcine pancreatic lipase inhibitory method was performed according to the protocol of Jaradat *et al*.[2019] [25]. With some modifications. A 500 µg/mL stock solution from each plant extract was dissolved in 10% dimethyl sulfoxide [DMSO], from which five different dilutions were prepared [50, 100, 200, 300 and 400 μg/ml]. Then, a 1 mg/ml stock solution of porcine pancreatic lipase was freshly prepared before use, which was dispersed in Tris-HCl buffer. The substrate used was *p*-nitrophenyl butyrate [PNPB] [Sigma-Aldrich, Germany], prepared by dissolving 20.9 mg in 2 mL of acetonitrile. In addition, for each working test tube, 0.1 ml of porcine pancreatic lipase [1 mg/ml] was mixed with 0.2 mL of the plant extract from each diluted solution series for each plant extract. The resulting mixture then completed to 1 ml by adding Tri-HCl solution and incubated at 37°C for 15 minutes. After this incubation period, 0.1 ml of p-nitrophenyl butyrate solution was added to each test tube. The mixture was then incubated for 30 minutes at 37°C. Pancreatic lipase activity determined by measuring the hydrolysis of PNPB into p-nitrophenolate ions at 410 nm using a UV-vis spectrophotometer; the same procedure was repeated for Orlistat [Sigma-Aldrich, Germany]. The inhibitory percentage of the anti-lipase activity was calculated using the following equation (2):

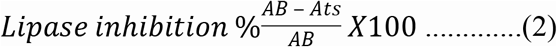

where AB is the recorded absorbance of the blank solution and Ats is the recorded absorbance of the tested sample solution.

### *In vitro* α-amylase inhibitory activity

The α-amylase inhibitory activity of *E. tirucalli* dried juice was carried out according to the standard method of [26] with minor modification. Briefly, each extract fraction was dissolved in few milliliters of 10% DMSO and then further dissolved in a buffer [[Na_2_HPO_4_/NaH_2_PO_4_ [0.02 M], NaCl [0.006 M] at pH 6.9] to give concentrations of 1000 μg/ml. The following dilutions were prepared [10, 50, 70, 100, 500 μg/ml]. A volume of 0.2 ml of porcine pancreatic α-amylase enzyme solution with concentration of [2 units/ml] was mixed with 0.2 ml of the plant juice then incubated for 10 min at 30°C. Thereafter, 0.2 ml a freshly prepared starch solution [1%] was added and the mixture was incubated for at least 3 min. The reaction was stopped by the addition of 0.2 ml Dinitrosalicylic acid [DNSA] then the mixture was diluted with 5 ml of distilled water and heated for 10 min in a water bath at 90°C. The mixture was left to cool down to room temperature, then the absorbance was taken at 540 nm. A blank was prepared following the same procedure replacing the plant fraction with 0.2ml of previous buffer.

Acarbose was used as positive control following the same procedure. The α-amylase inhibitory activity was calculated using the following equation:

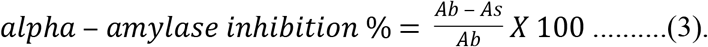

where:

A_b_: is the absorbance of blank

A_S_: is the absorbance of tested sample or control.

### α-glucosidase inhibitory activity

The α-glucosidase inhibitory activity of the dried *E. tirucalli* juice was carried out in accordance to the standard biochemical method with minor modification [27]. In each eppendorf tube a reaction mixture containing 50 μl phosphate buffer [100 mM, pH = 6. 8], 10 μl alpha-glucosidase [1 U/ml], and 20 μl of varying concentrations of the plant juice [100, 200, 300, 400 and 500 mg/ml] which was incubated at 37°C for 15 min. Then pre-incubated 20 μl of [5 mM] *p*-NPG was added as a substrate of the reaction and again incubated at 37°C for further 20 min. The reaction was terminated by adding 50 μl Na_2_ CO_3_ [0.1M]. The absorbance of the released *p*-nitrophenol was measured by a UV/Vis spectrophotometer at 405 nm. Acarbose, with similar concentrations to *E. tirucalli* dried juice, was used as a positive control.

Inhibition percentage was calculated using the following equation:

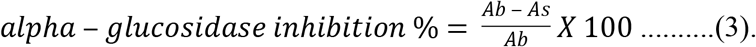

Where:

A_b_: is the absorbance of blank

A_S_: is the absorbance of tested sample or control.

### Cell proliferation assay for Caco-2 cells

Colorectal adenocarcinoma cells [Caco-2, ATCC® HTB-37™] were propagated in RPMI-1640 media, followed by the addition of 10% fetal bovine serum, also 1% penicillin-streptomycin antibiotics, and 1% l-glutamine. Caco-2 cells were grown in a moist atmosphere which contain 5% CO_2_ at 37°C. Cells were implanted at 2.6 × 10^4^ cells/well in a 96-well plate. After 24h, the cells were aggregated, and media were changed then cells were incubated with 5, 2.5, 1.25, 0.625, 0.3125 mg/ml for 24 h. Cell viability was defined by Cell Titer 96® Aqueous One Solution Cell Proliferation [MTS] Assay according to the manufacturer’s instructions [Promega Corporation, Madison, WI]. At the end of treatment, 20 μl of MTS solution/100 μl of media was added to every single well and were incubated for 2 h at 37°C. Finally, absorbance was measured at 490 nm.

### Antimicrobial method

#### Microbial strains

Reference microbial strains were obtained from American Type Culture Collection [ATCC]. Bacterial strains were *Staphylococcus aureus* [ATCC 25923], *Shigella sonnie* [ATCC 25931]*, Pseudomonas aeruginosa* [ATCC 27853]*, Escherichia coli* [ATCC 25922] and *Enterococcus faecium* [ATCC 700221]. While, the fungal strains were *Candida albicans* [ATCC 90028] and *Epidermophyton floccosum* [ATCC52066]. However, to carry out the antimicrobial activity, the *E. tirucalli* latex was dissolved in water to prepare 50 mg/ml solution.

### Micro plate broth dilution method

This method was carried out according to [28] for determination of minimal concentration inhibition [MIC] for both bacteria and yeast reference strains. Plant extract was examined in duplicate in each run. Using multichannel micropipette, 100 µl Mueller-Hinton II Broth [Becton Dickinson, France] was pipetted into each well of a 96 – well plate. Then in the first row a 100 µl from plant extract was placed and mixed thoroughly, followed by transferring a 100 µl to next raw. This was repeated to the 10^th^ row then the 11^th^ row has got 100 µl plant extract representing the negative control for bacterial growth. The 12^th^ row in the plate was left as positive control for microbial growth containing no plant extract. After this serial dilution, the examined plant extract concentrations were [µg/µl]; 50, 25, 12.5, 6.25, 3.125, 1.562, 0.781, 0.390 etc.

Bacterial suspension from fresh culture was prepared and turbidity was made equivalent to that of 0.5 McFarland standard so that the concentration of bacterial isolate was about 1.5× 10^8^ CFU/ml. Then the bacterial suspension was diluted 1:3 in sterile broth. Bacterial suspension [1µl] was mixed thoroughly with solutions in rows 1-10 and 12 while leaving raw 11 with no bacterial suspension. The plates were incubated at 35 °C for 18 hours.

For yeast, a yeast suspension was prepared with turbidity equivalent to 0.5 McFarland standard. The yeast suspension was diluted 1:1000 in sterile broth. A 100µl was transferred to a 96-well plate as described above. The plate was incubated at 35°C for 48 hours.

### Agar diffusion well-variant method

This method was made according to [29, 30]. Bacterial suspension with turbidity equivalent to 0.5 McFarland standard was prepared from fresh culture. A sterile cotton swab was used to distribute the bacterial inoculum uniformly on surface of Mueller-Hinton II agar. In each inoculated plate using sterile glass cylinder, 5 of 6-mm diameter wells were made with 2.5-cm gaps. In each well, 80 µl plant extract were placed. The plates were then incubated at 37 ºC for 18 hours. The diameters of inhibition zones were measured. Each plant extract test was made in duplicate and the average of diameter was calculated. Ciprofloxacin antibiotic [50 mg/ml] was used as positive control.

### Agar dilution method

#### Preparation of Sabouraud dextrose agar [SDA]

Twelve tubes after autoclaving were containing 1 ml SDA. They were divided into 2 groups and numbered with E1-E6. Tubes, then, were placed in the water bath at 45 ºC to remain in a liquid state. In addition, tubes containing microfiltered plant extracts were also placed in the same water bath.

One ml of the extract was transferred to E1 then 1 ml serially transferred to tubes E2-E5 respectively. All tubes were positioned in slant to solidify. Tube E6 was left as a control. The same steps were repeated on the second set of tubes.

#### Yeast Enumeration

Yeast cultures were grown on potato dextrose broth and transferred every 72 h to assure purity. A freshly prepared yeast culture grown on potato dextrose agar was flooded with10 ml of 0.05% Tween 20 in normal saline, then 2ml of the yeast suspension was pipetted twice by a micropipette and transferred to a tube containing 5 ml normal saline. The yeast suspension was prepared to achieve a turbidity equivalent to 0.5 McFarland.

Twenty-µl of suspension of fungus was aseptically transferred to each tube that previously numbered from E1-E6 of each group except for tubes numbered E5.

The tubes were left for one day then incubated at 25°C for further 14 days.

## Results

### Digestive enzymes inhibitory activity

The inhibitory activities against α-amylase, α-glucosidase and lipase enzymes were conducted according to standard biochemical tests. The results showed that the dried juice of *E. tirucalli* plant has a strong α-glucosidase and lipase enzymes inhibitory activity with IC_50_ values of 79.43±0.38 and 39.8±0.22, respectively as shown in Table 1 and Fig. 2. While the inhibitory activity against α-amylase was weak in comparison with Acarbose drug.

**Table 1.** The α-glucosidase inhibitory activity and IC_50_ values of *E. tirucalli* and Acarbose.

| Conc. | Acarbose | *E. tirucalli* |
| --- | --- | --- |
| 0 | 0±0 | 0±0 |
| 100 | 65.8±0.42 | 57.76±0.47 |
| 200 | 67.75±0.35 | 58.1±0.95 |
| 300 | 73.2±0.42 | 65.7±0.23 |
| 400 | 85.35±0.35 | 65.7±0.23 |
| 500 | 92.22±0.11 | 66.8±0 |
| α-glucosidase<br>inhibitory<br>activity IC<sub>50</sub><br>[μg/ml], ±SD | 37.1 ±0.33 | 79.43±0.38 |

**Figure 1.**
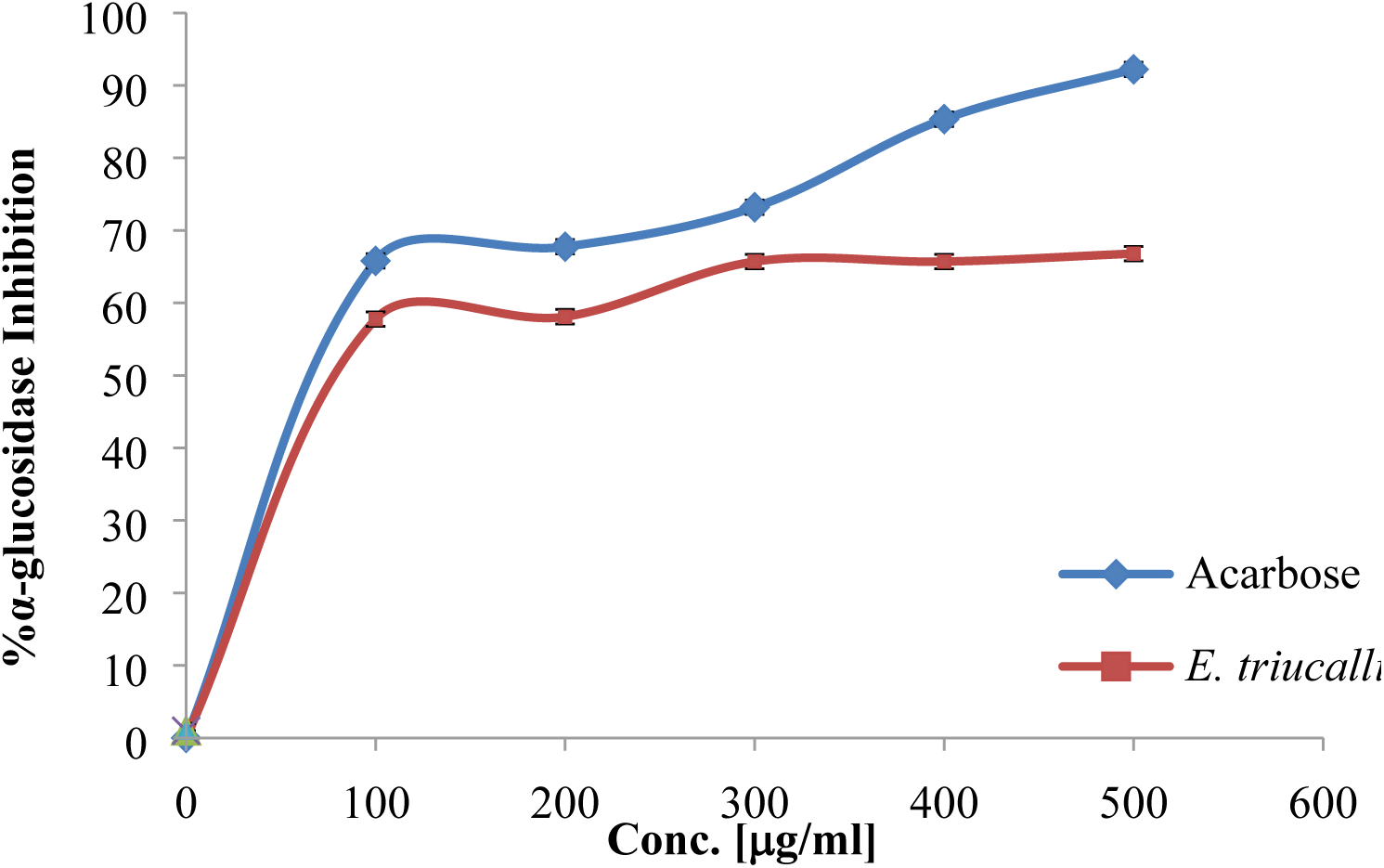
The dried juice of *E. tirucalli* plant and Acarbose antidiabetic drug α-glucosidase inhibitory activities

**Figure 2.**
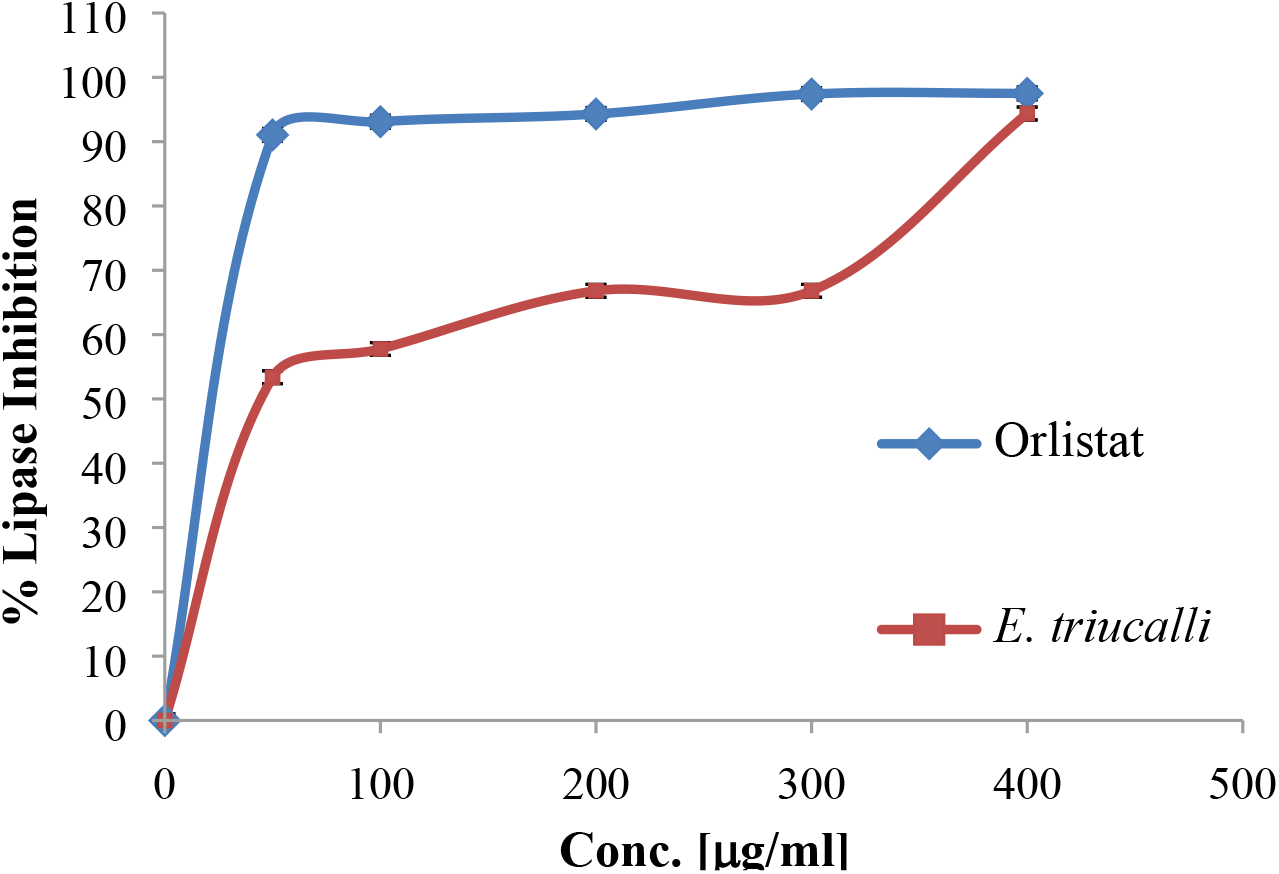
The dried juice of *E. tirucalli* plant and Acarbose antidiabetic drug lipase inhibitory activities. A 500 µg/ml stock solution from each plant extract was dissolved in 10% dimethyl sulfoxide (DMSO), The substrate used was *p*-nitrophenyl butyrate (PNPB) and 0.1 ml of porcine pancreatic lipase (1 mg/ml) was mixed

### Antioxidant activity

The DPPH method was utilized to assess the antioxidant potential of the dried juice of *E. tirucalli* plant and Trolox. Table 4 and Fig. 4 depicted that the plant has mild antioxidant potentials in comparison to Trolox.

**Table 2.**
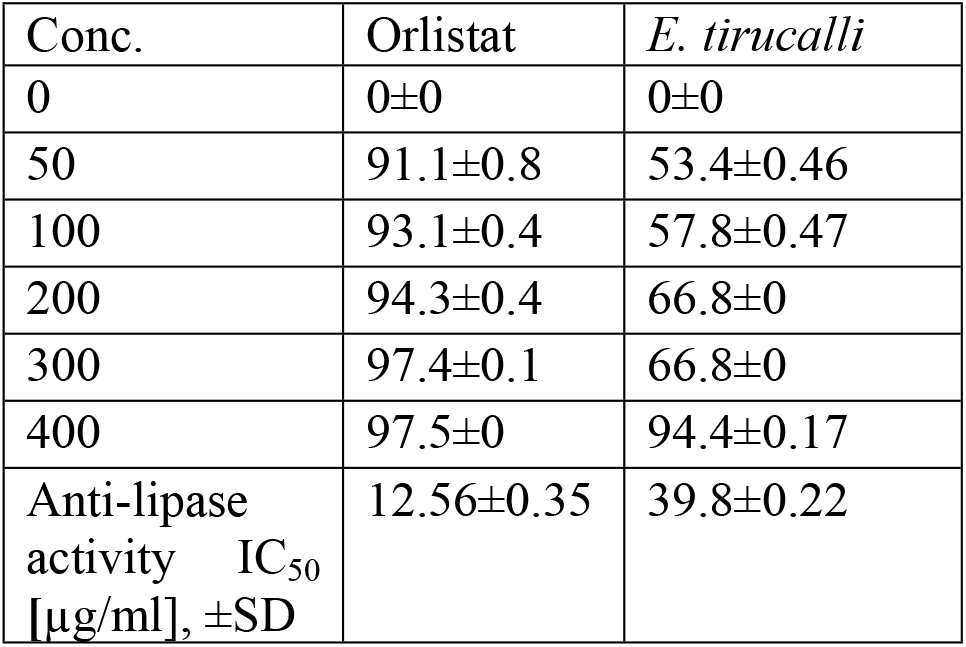
Anti-lipase inhibitory activity and IC_50_ values of *E. tirucalli* and Orlistat.

**Table 3.**
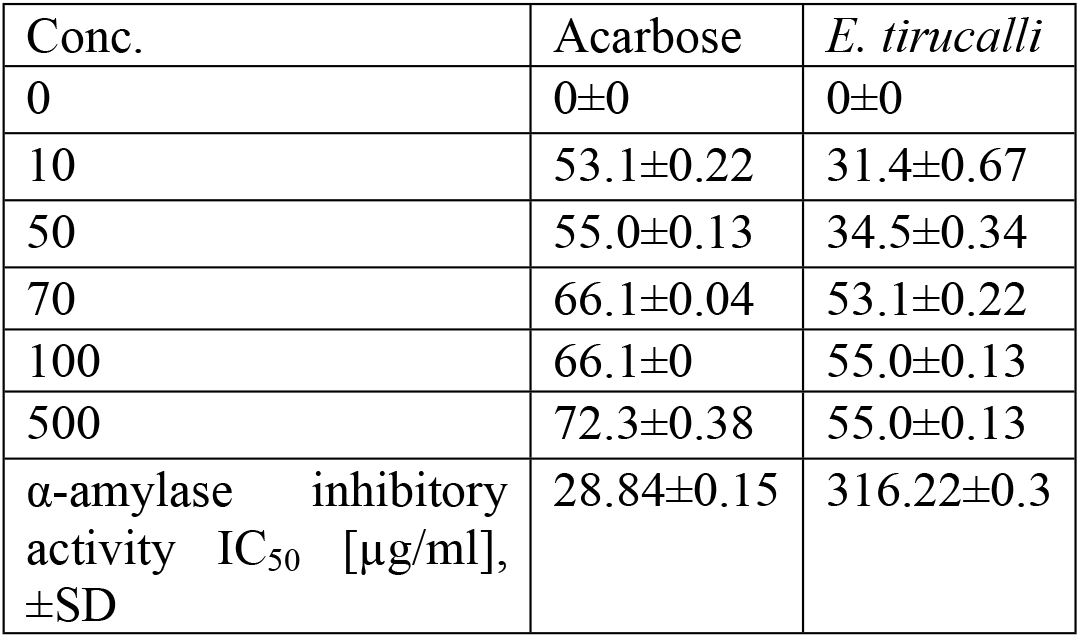
The α-amylase inhibitory activity and IC_50_ values of *E. tirucalli* and Acarbose

| Conc. | Acarbose | *E. tirucalli* |
| --- | --- | --- |
| 0 | 0±0 | 0±0 |
| 10 | 53.1±0.22 | 31.4±0.67 |
| 50 | 55.0±0.13 | 34.5±0.34 |
| 70 | 66.1±0.04 | 53.1±0.22 |
| 100 | 66.1±0 | 55.0±0.13 |
| 500 | 72.3±0.38 | 55.0±0.13 |
| α-amylase inhibitory activity IC<sub>50</sub> [μg/ml], ±SD | 28.84±0.15 | 316.22±0.3 |

**Table 4.**
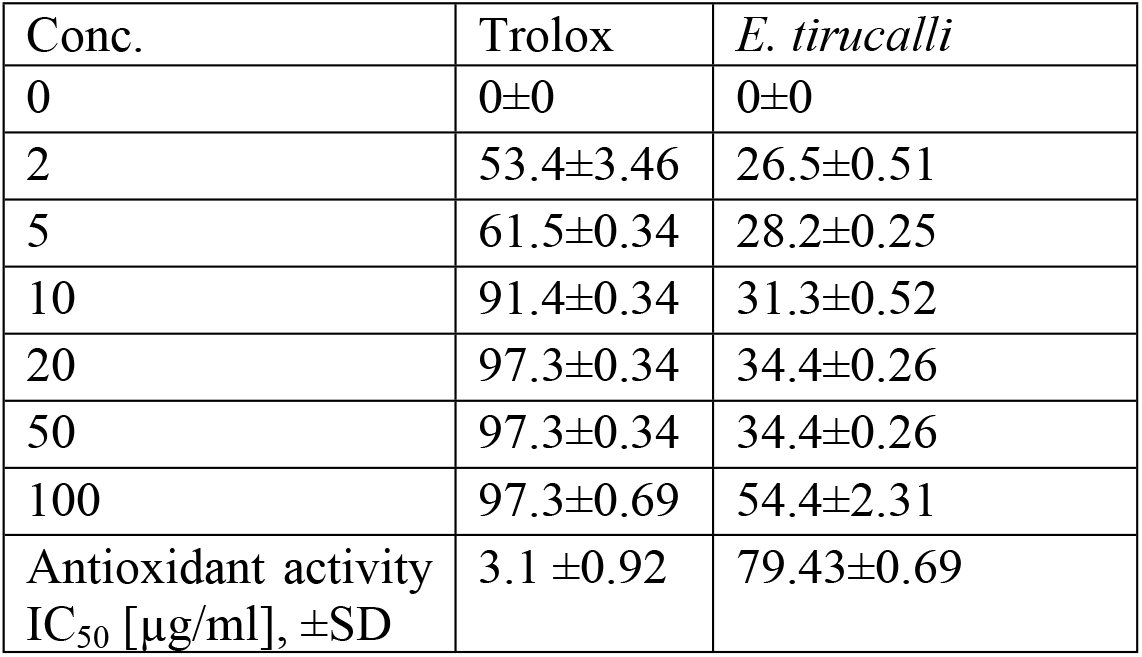
DPPH free radical scavenging property and IC_50_ values of *E. tirucalli* and Trolox.

| Conc. | Trolox | *E. tirucalli* |
| --- | --- | --- |
| 0 | 0±0 | 0±0 |
| 2 | 53.4±3.46 | 26.5±0.51 |
| 5 | 61.5±0.34 | 28.2±0.25 |
| 10 | 91.4±0.34 | 31.3±0.52 |
| 20 | 97.3±0.34 | 34.4±0.26 |
| 50 | 97.3±0.34 | 34.4±0.26 |
| 100 | 97.3±0.69 | 54.4±2.31 |
| Antioxidant activity<br>IC<sub>50</sub> [µg/ml], ±SD | 3.1 ±0.92 | 79.43±0.69 |

**Figure 3.**
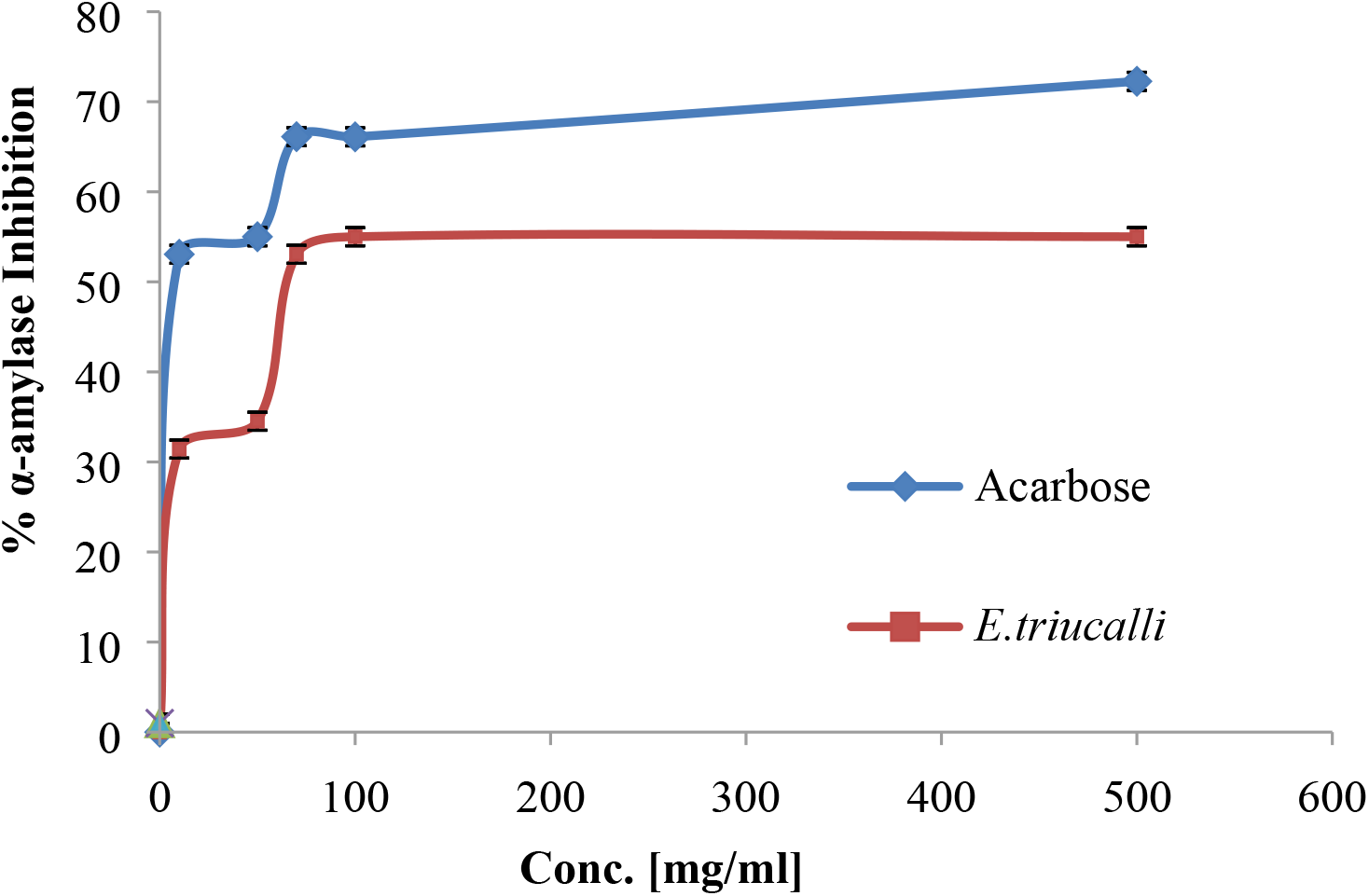
The dried juice of *E. tirucalli* plant and Acarbose antidiabetic drug α-amylase inhibitory activities

**Figure 4.**
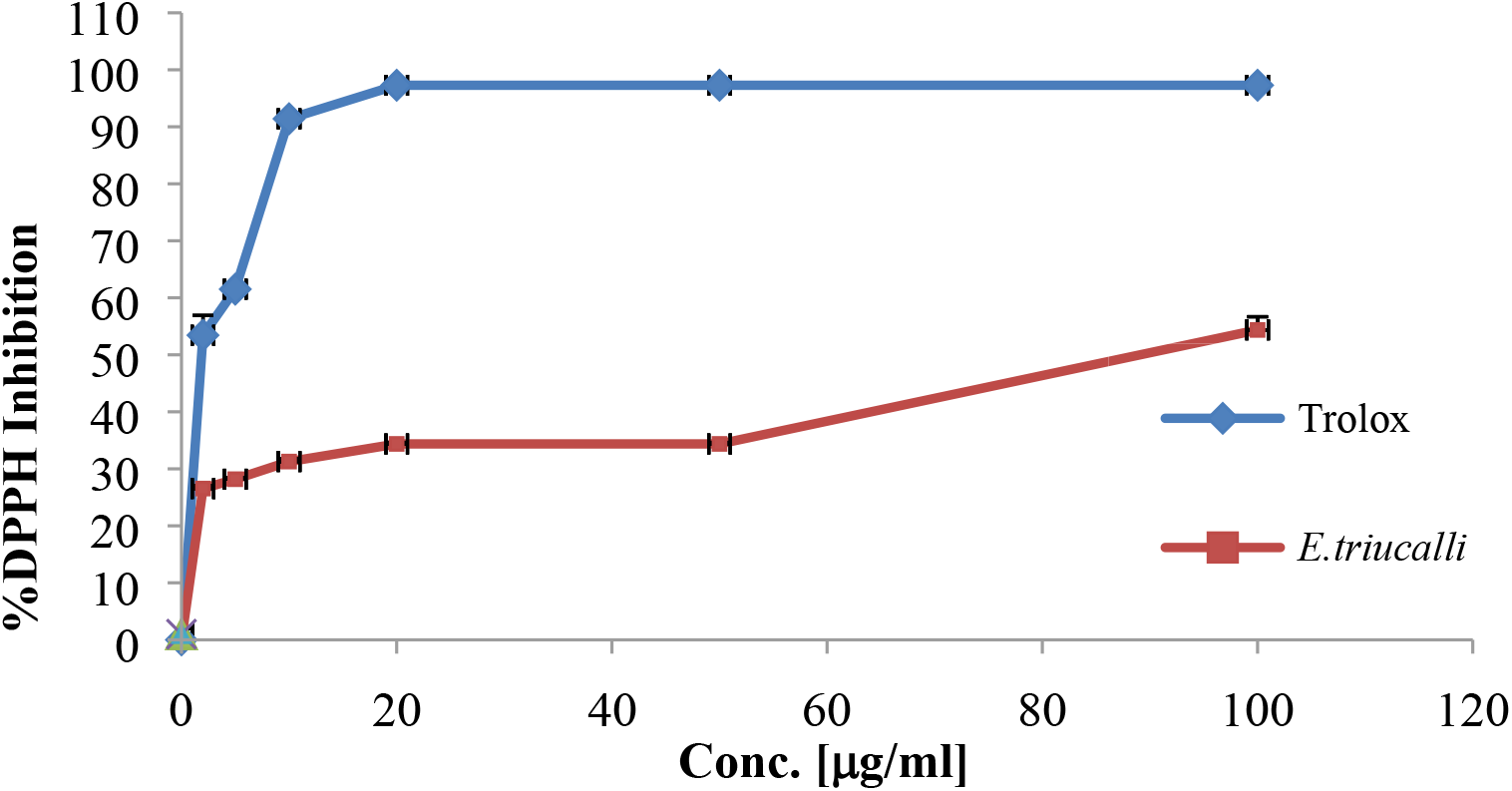
DPPH free radical scavenging property of *E. tirucalli* and Trolox. Methanolic stock solution (lmg/ml) was set for *E. tirucalli* dried juice. Plant working solution (1 ml) was mixed with 1ml freshly prepared DPPH (0.002 g/ml) methanolic solution. The solutions were incubated at room temperature (25°C) in a dark place for 30 min. Then, their optical densities were measured by the UV/Vis spectrophotometer at 517 nm.

### Agar diffusion well-variant method

The juice of *E. tirucalli* plant was milky and even after dilution all wells remained unclear, so micro broth dilution method was not suitable for evaluation of antimicrobial activities. There wasn’t any inhibition zone around wells. On showed inhibition zones, which were 4, 5.5, 5.5, 5.5 and 5.8 cm for *S. aureus, Shigella sonnie, Pseudomonas aeruginosa, E. coli* and *E. faecium*, respectively.

### Agar dilution method

Anti-mold effects of the plant extract are shown on the Table 6. There was no growth of the fungus in the tubes numbered from E1-E5, but fungus growth was observed in the tube E6. These results revealed that the plant dried juice inhibited the growth of *Epidermophyton floccosum*.

**Table 5.**
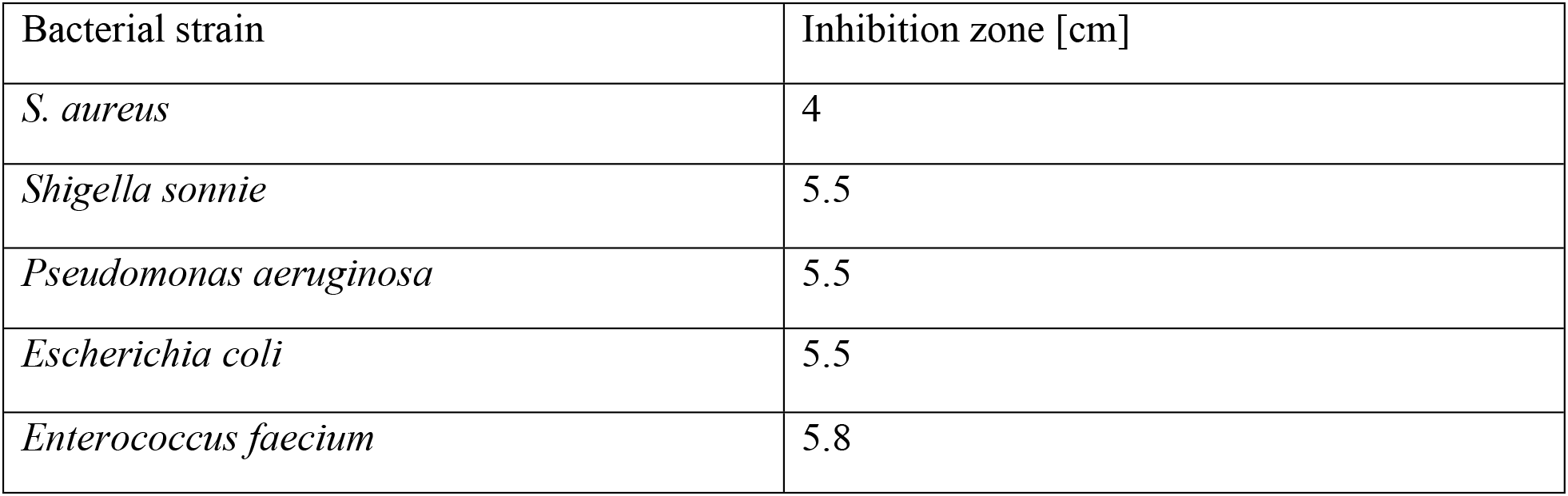
Inhibition zone of Ciprofloxacin [positive control]

**Table 6.** The growth of the fungus on the slant agar.

| Tube number | Growth result |
| --- | --- |
| E1 | No fungus growth |
| E2 | No fungus growth |
| E3 | No fungus growth |
| E4 | No fungus growth |
| E5 | No fungus growth |
| E6 | Fungus growth |

### Cytotoxic effect of extracts derived from *Euphorbia tirucalli*

As shown in Fig. 5 treatment of Caco-2 cells with 5, 2.5, 1.25, 0.625, 0.3125 mg/ml of *Euphorbia tirucalli* dried juice induced cytotoxicity significantly [p≤0.001] by approximately 86%, 88%, 80%, 37% and 13% respectively.

**Figure 5.**
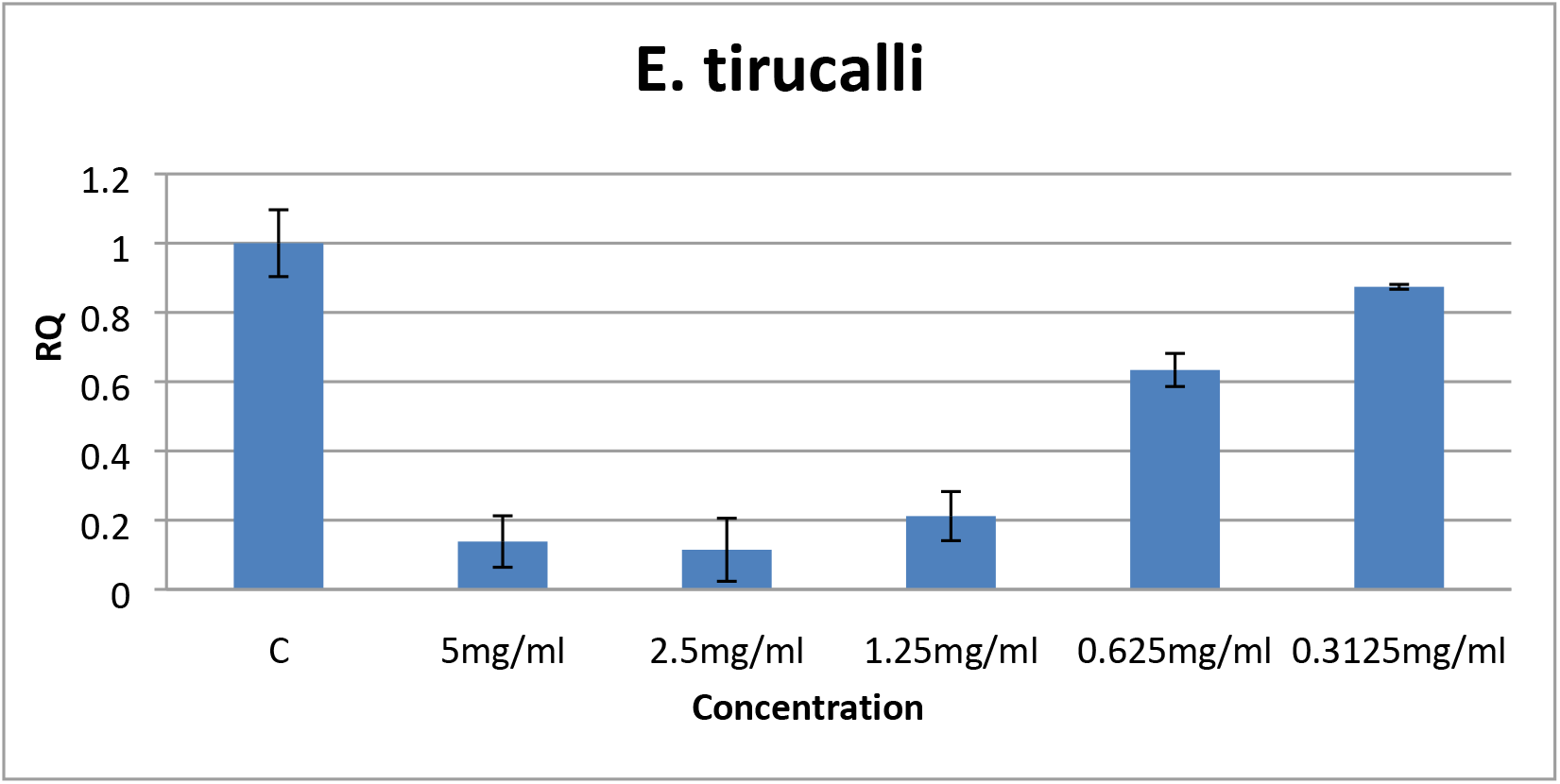
The cytotoxicity of *E. tirucalli* on the Caco-2 cells. The effect of *E. tirucalli* on the cytotoxicity of Caco-2 cells. Caco-2 cells were treated with 5, 2.5, 1.25, 0.625, 0.3125 mg/ml of *Euphorbia tirucalli* crude extract for 24 h. Cytotoxicity was determined by MTT assay. Results were depicted as relative quantities [RQ] compared to the control [with only media; C]. \**P<0.001*. Error bars represent SD.

## Discussion

Plants are natural source of many bioactive compounds. Some plants exhibit a wide range of bioactivities due to production of secondary metabolites they produce. Such metabolites are usually produced to protect the plants from the harsh environment and natural enemies. In this context *E. tirucalli*, though an ever-green plant, has been rarely fed by herbivores. Moreover, reports showed that it has few pests and diseases due to its poisonous latex. In this study we examined the bioactivities of the latex of *E. tirucalli.* The chemical composition of the latex was reported in many researches. Its complexity may explain the wide varieties of the functions it has. The extract activity against digestive enzymes was a half and a third of that of the controls for glucosidase and lipase respectively while it was weaker with amylase this may suggest that the extract compete with the substrates on the active sites, however, in case of amylase which reacts with larger molecule such as starch with many possible sites of reaction this will make competition process less likely. Moreover, the variety of the extract constituents may result in different affinities to react with different proteins [enzymes]. This is in agreement with many researches which found that plant extracts have variable activities against digestive enzymes due to levels of polyphenols, flavonoids, terpenoids etc [31,32]. On the other hand, the current study of the antimicrobial activity of the crude alcoholic extracts of *E. tirucalli* against some types of pathogens has shown that it exhibited a low MIC against *S. aureus* and other bacteria under investigation. This has a great significance in the healthcare delivery system, so it could be used as an alternative to orthodox antibiotics in treatment of some infections caused by these microbes, especially as they usually developing resistance to the known antibiotics.

Antimicrobial activity can be explained due to the fact that *E. tirucalli* is high in tannins. Tannins are known to be surfactants and may inactivate microbial adhesions and also complex with membrane polysaccharides. Many plant genetic sources have been analyzed for their active components have antimicrobial activities. The antimicrobial activity exhibited by *E. tirucalli* may be attributed to the various active constituent’s present, which also because of their individual or combined action, shows antimicrobial activity [33, 34]. Moreover, E. tirucalli showed complete inhibition of the yeast cells used in this experiment.

Antioxidant ability of bioactive compounds is very crucial in fighting against tumors as it was found that free radicals are significant players in the etiology of cancer. *E. tirucalli’s* extract was found to have mild ability in reacting with DPPH compared with control. This is may be due to that the extract was mechanically prepared and was mixed with methanol, meaning that not all active constituents were presented in the extract. Polyphenols could play an important role in this context and their effects have been studied *in vivo* and *in vitro*. Many polyphenols, such as anthocyanin, proanthocyanins, flavonoid, resveratrol, tannins, epigallocatechin-3-gallate, and gallic acid, have been tested; all of them showed promising effects however, their mechanisms of action were variable [35]. *E tirucalli’s* extract showed potential toxicity against Caco-2 cell line. A lot of pharmacological activities of E. tirucalli have been documented by many studies as molluscicidal activity, antimicrobial activity, antiherpetic and anti-mutagenic activity. The latex also shows anti-carcinogenic activities. In the northeast region of Brazil, the extract of *E. tirucalli* is used; as an antibacterial agent; a laxative effect; to treat intestinal parasites; to treat cough, asthma, rheumatism, cancer, sarcoma, and epithelioma skin tumors [36].

## Conclusion

Taken together, our results suggest that the dried latex of *E. tirucalli* induces cytotoxicity to human cancer cells. However, the exact mechanism of action of the plant on the examined cells needs to be further explored. Moreover, the dried juice of *E. tirucalli* plant has a strong α-glucosidase and lipase enzymes inhibitory activity. While the inhibitory activities against α-amylase was weak in comparison with Acarbose drug. In addition, the *E. tirucalli* plant dried juice showed mild antioxidant effect. All these facts suggest the benefits of the *E. tirucalli* plant use in the pharmaceutical and functional foods industries due to its potential health benefits.

## Conflict of interest

The authors declare that they have no conflict of interest with this research or its outcomes.

## Acknowledgment

The authors would like to thank the instructors at the Department of Pharmacy for their support and efforts.

## Fund

No source of fund.

---

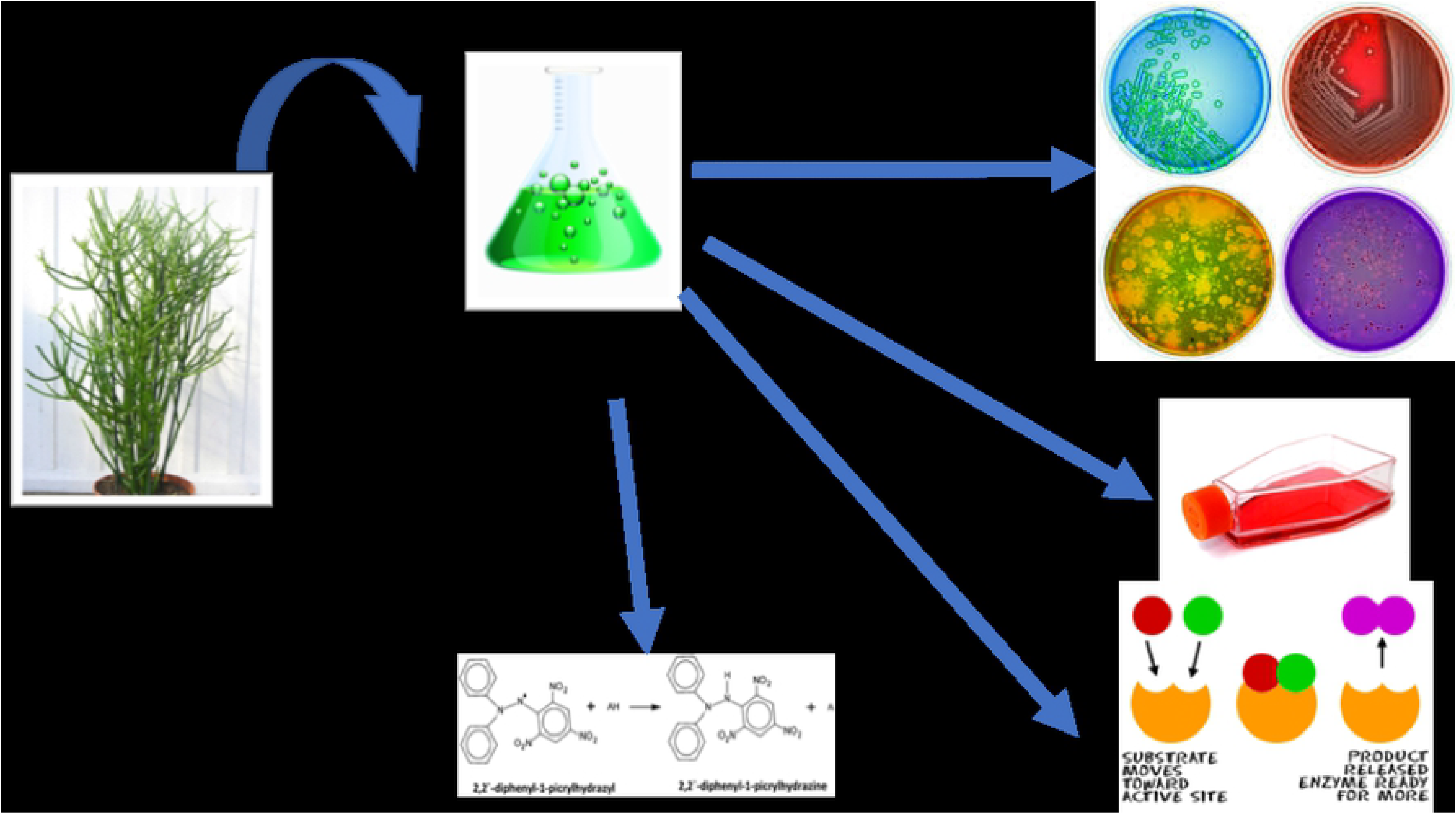

## References

1. Ebrahimi A, Atashi A, Soleimani M, Mashhadikhan M, Kaviani S. Comparison of anticancer effect of Pleurotus ostreatus extract with doxorubicin hydrochloride alone and plus thermotherapy on erythroleukemia cell line. Journal of Complementary and Integrative Medicine. 2017;15(2):doi: 10.1515/jcim-2016-0136.

2. Khalil A. Role of Biotechnology in Alkaloids Production. Catharanthus roseus. Germany: Springer; 2017; 59–70.

3. Head GA. Cardiovascular and metabolic consequences of obesity. Frontiers in physiology. 2015;6:32.

4. Tazelaar HD, Lilenbaum RC. Pathology of lung malignancies. Up To Date, Nicholson A, Jett J, Lilenbaum R [Section Editors], Up To Date, Vora S[consultado 18 Mar 2017] Disponible en: https://uptodate_publicaciones_saludcastillayleones/contents/pathology-of-lung-malignancies.1-19.

5. Mokdad AH, Ford ES, Bowman BA, Dietz WH, Vinicor F, Bales VS, et al. Prevalence of obesity, diabetes, and obesity-related health risk factors, 2001. Jama. 2003;289(1):76–9.

6. World Health Organization. Cancer WHO; 2019 [cited 2019 23, April]. Available from: https://www.who.int/news-room/fact-sheets/detail/cancer.

7. McAdam AJ, Hooper DC, DeMaria A, Limbago BM, O’Brien TF, McCaughey B. Antibiotic resistance: how serious is the problem, and what can be done? Clinical chemistry. 2012;58(8):1182–6.

8. Valko M, Jomova K, Rhodes CJ, Kuča K, Musilek K. Redox-and non-redox-metal-induced formation of free radicals and their role in human disease. Archives of toxicology. 2016;90(1):1–37.

9. Goszcz K, Duthie GG, Stewart D, Leslie SJ, Megson IL. Bioactive polyphenols and cardiovascular disease: chemical antagonists, pharmacological agents or xenobiotics that drive an adaptive response? British journal of pharmacology. 2017;174(11):1209–25.

10. Khan AQ, Ahmed Z, Malik A, Afza N. The structure and absolute configuration of cyclotirucanenol, a new triterpene from *Euphorbia tirucalli* Linn. Zeitschrift für Naturforschung B. 1988;43(8):1059–62.

11. Uchida H, Ohyama K, Suzuki M, Yamashita H, Muranaka T, Ohyama K. Triterpenoid levels are reduced during *Euphorbia tirucalli* L. callus formation. Plant biotechnology. 2010;27(1):105–9.

12. Khan AQ, Rasheed T, Malik A. Tirucalicine: a new macrocyclic diterpene from *Euphorbia tirucalli*. Heterocycles. 1988;27(12):2851–6.

13. Gupta N, Vishnoi G, Wal A, Wal P. Medicinal value of *Euphorbia tirucalli*. Systematic Reviews in Pharmacy. 2013;4(1):40–6.

14. Nielsen PE, Nishimura H, Liang Y, Calvin M. Steroids from Euphorbia and other latex-bearing plants. Phytochemistry. 1979;18(1):103–4.

15. Waczuk EP, Kamdem JP, Abolaji AO, Meinerz DF, Bueno DC, do Nascimento Gonzaga TKS, et al. *Euphorbia tirucalli* aqueous extract induces cytotoxicity, genotoxicity and changes in antioxidant gene expression in human leukocytes. Toxicology Research. 2015;4(3):739–48.

16. Schmelzer G. In Schmelzer GH, Gurib-Fakim A,[Eds.], Arroo R,[Associate Ed.] Lemmens RHMJ, Oyen LPA,(General Ed.). Plant Resources of Tropical Africa: Medicinal plants. 2008;11:1.

17. Duke JA. Handbook of energy crops: Springer; 1983. https://www.hort.purdue.edu/newcrop/duke_energy/Euphorbia_tirucalli.html

18. Siddiqui M, Alam M, Trivedi P. In Nematode Management in Plants. India: Scientific Publishers,; 2003.

19. Mwine TJ, Van Damme P. *Euphorbia tirucalli* L.(Euphorbiaceae): the miracle tree: current status of available knowledge. Scientific Research and Essays. 2011;6[23]:4905–14.

20. Jurberg P, Cabral Neto JB, Schall VT. Molluscicide activity of the” Avelós” plant (*Euphorbia tirucalli,* L.) on Biomphalaria glabrata, the mollusc vector of schistosomiasis. Memórias do Instituto Oswaldo Cruz. 1985;80(4):423–7.

21. Yadav R, Srivastava V, Chandra R, Singh A. Larvicidal activity of latex and stem bark of *Euphorbia tirucalli* plant on the mosquito *Culex quinquefasciatus*. The Journal of communicable diseases. 2002;34(4):264–9.

22. Madureira AM, Ferreira M-JU, Gyémánt N, Ugocsai K, Ascenso JR, Abreu PM, et al. Rearranged jatrophane-type diterpenes from euphorbia species. Evaluation of their effects on the reversal of multidrug resistance. Planta medica. 2004;70(10):45–9.

23. Bani S, Kaul A, Khan B, Gupta VK, Satti NK, Suri KA, et al. Anti-arthritic activity of a biopolymeric fraction from *Euphorbia tirucalli*. Journal of ethnopharmacology. 2007;110(1):92–8.

24. Imai S, Sugiura M, Mizuno F, Ohigashi H, Koshimizu K, Chiba S, et al. African Burkitt’s lymphoma: a plant, *Euphorbia tirucalli*, reduces Epstein-Barr virus-specific cellular immunity. Anticancer research. 1994;14(3A):933–6.

25. Jaradat, N. A., Al-lahham Saad, Zaid, A. N., Hussein, F., Issa, L., Abualhasan, M. N. et al. *Carlina curetum* plant phytoconstituents, enzymes inhibitory and cytotoxic activity on cervical epithelial carcinoma and colon cancer cell lines. European Journal of Integrative Medicine, 100933. doi:10.1016/j.eujim.2019.100933

26. Nyambe-Silavwe H, Villa-Rodriguez JA, Ifie I, Holmes M, Aydin E, Jensen JM, et al. Inhibition of human α-amylase by dietary polyphenols. Journal of Functional Foods. 2015;19:723–32.

27. Ademiluyi AO, Oboh G. Soybean phenolic-rich extracts inhibit key-enzymes linked to type 2 diabetes [α-amylase and α-glucosidase] and hypertension [angiotensin I converting enzyme] in vitro. Experimental and Toxicologic Pathology. 2013;65(3):305–9.

28. Forbes BA, Sahm DF, Weissfeld AS Laboratory methods and strategies for antimicrobial susceptibility testing. St. Louis: Mosby; 2007.

29. Falahati M, Omidi Tabrizib N, Jahaniani F. Anti dermatophyte activities of *Eucalyptus camaldulensis* in comparison with Griseofulvin. Iranian Journal of Pharmacology and Therapeutics. 2005;4(2):80–0.

30. BA Forbes, DF Sahm, Weissfeld A. Study Guide for Bailey and Scott’s Diagnostic Microbiology. UK: Elsevier Health Sciences; 2016.

31. Yousaf AM, Susoma J, Ah JH, Oh JH, Young CH, Sue CJ. Coumarins from angilica decursiva inhibit-glucosidase activity and protein tyrosinane phosphatase 1B. Chemico-Biological Interactions. 2016; 252 (25). 93–101.

32. Zhang, AJ.; Rimando AM.; Mizuno CS.; Mathews ST. Alph-glucosidase inhibitory effect of resveratrol and piceatannol. The Journal of Nutritional Biochmestry. 2017; 47. 86–93.

33. Suphachai Charoensin. Antioxidant and anticancer activities of *Moringa oleifera* leaves. Journal of Medicinal Plant Research. 2014; 8(7), 318–3257

34. Mahesh B. and Satish S.Antimicrobial Activity of Some Important Medicinal Plant Against Plant and Human Pathogens World Journal of Agricultural Sciences. 20084 (S): 839–843.

35. Jiménez S, Gascón S, Luquin A, Laguna M, Ancin-Azpilicueta C, Rodríguez-Yoldi MJ Rosa canina Extracts Have Antiproliferative and Antioxidant Effects on Caco-2 Human Colon Cancer. PLoS ONE. 2016; 11(7): e0159136. doi:10.1371/journal.pone.0159136

36. Upadhyay B, Singh KP, Ashwani K. Ethno-Medicinal, Phytochemical and Antimicrobial Studies of *Euphorbia tirucalli* L. Journal of Phytology. 2010;10:110–6.

